# A balanced CAR-T cell metabolism driven by CD28 or 4-1BB co-stimulation correlates with clinical success in lymphoma patients

**DOI:** 10.1101/2024.09.27.615167

**Authors:** Molly S. Cook, Katie R. Flaherty, Sophie Papa, Reuben Benjamin, Anna Schurich

## Abstract

Chimeric antigen receptor (CAR) T cell therapy has led to unprecedented success in treating relapsed/refractory diffuse large B cell lymphoma (DLBCL). The most common CAR-T cell products currently in the clinic for DLBCL differ in their co-stimulation moiety, containing either CD28 or 4-1BB, which initiate distinct signalling pathways. Previous work has highlighted the importance of T cell metabolism in fuelling anti-cancer function. We have studied the metabolic characteristics induced by CD28 versus 4-1BB co-stimulation in patient CAR-T cells *ex vivo*. Our data show that in patients, CD28 and 4-1BB drive significantly divergent metabolic profiles. CD28 signalling endows T cells with a preferentially glycolytic metabolism supporting an effector phenotype and increased expansion capacity, while 4-1BB co-stimulation preserves mitochondrial fitness and results in memory-like differentiation. Despite this divergent programming, T cells in patients responding successfully to therapy were metabolically similar, irrespective of co-stimulator, suggesting that efficient fuelling of CAR-T cells in lymphoma requires a ‘*balanced’* metabolism. In contrast, CAR-T cells in non-responders were pushed to their metabolic extremes.

**One sentence summary:** CD28 and 4-1BB signalling drive divergent metabolic profiles in patient CAR-T cells, however patients that respond to therapy have CAR-T cells that maintain metabolic balance.

## Introduction

CAR-T cell therapy has shown great success in the treatment of relapse/refractory DLBCL, the most common form of lymphoma. Despite this, around 50% of patients ultimately experience therapeutic failure^1^. Clinically approved CD19-targeting CAR-T cells contain distinct co-stimulators, either CD28 or 4-1BB (CD137). It was the inclusion of these co-stimulation domains into CAR constructs that paved the way for successful clinical application^2–4^. T cell activation, effector functions and memory formation require specialised metabolic programmes, which are modulated by co-stimulatory signalling. CD28, a member of the immunoglobulin superfamily of receptors, is a key co-stimulator of T cell activation. CD28 augments metabolic reprogramming via Pi3K/Akt/mTORC1 signalling, promoting an increased glycolytic metabolism^5,6^. Glycolysis supports biosynthesis of molecules required for rapid proliferation and effector functions. In contrast, signalling via 4-1BB, a member of the TNFR superfamily, promotes increased mitochondrial oxidative phosphorylation and central memory differentiation^7,8^. Increased mitochondrial biogenesis and respiratory capacity are linked to enhanced anti-cancer responses in various models^7,8^ and 4-1BB co-stimulated CAR-T cells have persisted in patients for a decade^9^. Despite this, a large clinical cohort study found superior clinical response provided by CD28 co-stimulated CAR-T cells in patients with DLBCL^10^. Studying metabolic profiles of CAR-T cells in patients has been challenging, often due to low CAR-T cell numbers in circulation or tissue. To address this lack of knowledge, we have made use of recent technical advances to compare CD28 and 4-1BB co-stimulated patient-derived CAR-T cells *ex-vivo* on a single cell level^11,12^. Our data show that CD28 co-stimulation drives increased glycolytic dependence supporting an ‘effector’ T cell profile. 4-1BB in contrast stimulates mitochondrial fitness and metabolism, and memory-like features. Of interest, in patients who achieved a complete response at 6 months, CAR-T cell metabolic profiles showed a ‘balanced’ phenotype with both glycolytic and mitochondrial capacity. In contrast CAR-T cells in non-responders displayed extreme polarisation towards either very high glycolysis in CD28 co-stimulated or suppressed glycolysis in 4-1BB co-stimulated T cells. Our data suggest that for successful response of CAR-T cells in DLBCL neither extreme metabolic feature is favourable. A deeper understanding of the metabolic trajectory supported by distinct CAR-constructs could lead to improved CAR-T therapies and better personalisation of CAR-T products in the future.

## Results

### CD28 and 4-1BB signalling differentially regulate the expression of glucose metabolism-associated proteins in patient CAR-T cells

To assess the impact of co-stimulation on CAR-T metabolism in patients, we analysed *ex vivo* CD19-targeting CAR-T cells from a total of 23 DLBCL patients, 14 patients who received a CD28-costimulated CAR-T product (axicabtagene ciloleucel) and 9 patients who received a 4-1BB-costimulated CAR-T product (tisagenlecleucel) (table 1). We could detect circulating live T cells expressing CAR on their surface by staining with an antibody against the single chain variable fragment, FMC63 (fig. 1 a). Upon activation, T cells upregulate glucose transporter 1 (GLUT-1) to support the increased use of glycolysis^13^. We therefore first stained for GLUT-1 expression in *ex vivo* CAR-T cells, and in line with the known signalling function of CD28 supporting glycolysis ^5,6^, we found that CD28-CAR-T expressed significantly higher levels of GLUT-1 (fig. 1b). Increased glycolysis is regulated by the transcription factor mTORc1^14^. We found CD28 CAR-T to have higher phosphorylation levels of mTOR at Ser2448 (fig. 1c). Additionally, CD28 co-stimulation also led to increased expression of the large neutral amino-acid transporter CD98 (fig. 1d), which supports T cell expansion^15^. Whilst CD28 co-stimulation has been associated with increased glycolysis, 4-1BB co-stimulation has been linked to an increase in mitochondrial function and a memory-like T cell phenotype^7^. We therefore stained for PGC1a, a transcriptional co-activator which has a known role in inducing mitochondrial biogenesis^8,16–18^. However, we did not find any significant difference in PGC1a expression between CD28 and 4-1BB CAR-T (fig. 1e). We also did not find a significant difference in the expression of CD71, the transferrin-receptor 1 transporter protein. CD71 is upregulated upon T cell activation and allows increased capacity to take up iron which can be utilized, for example, in enzymes of the mitochondrial electron transport chain^19^ (fig. 1f). Mitochondrial respiration can be fueled through fatty acid oxidation. We assessed expression of CPT1A (carnitine palmitoyl transferase 1 alpha), an enzyme regulating fatty acid oxidation by shuttling long-chain fatty acids into the mitochondria, but found no significant difference between CD28-CAR-T and 4-1BB-CAR-T (fig. 1g). Our data suggest that specifically the expression of glycolysis-associated proteins is differentially regulated by CD28 and 4-1BB in patient CAR-T cells.

**Table 1.** Patient Information. Prior disease stage determine by FDG-PET scan and ranked from 1 to 5 (low to heavy disease burden) using the Deauville scale. * Mean shown with range in brackets.

|  | Patients treated with CD28-CAR-T | Patients treated with 4-1BB-CAR-T |
| --- | --- | --- |
| No. Patients | 14 | 9 |
| Sex | 11M: 3F | 6M: 3F |
| Age* | 57.5 (38 - 72) | 69.5 (53 - 78) |
| BMI* | 27.3 (22.6 - 36.3) | 25.8 (18.5 - 33.3) |
| Prior disease stage* | 4.4 (2 - 5) | 4.4 (2 - 5) |
| 6 month survival (%) | 80 | 70 |

**Figure 1.**
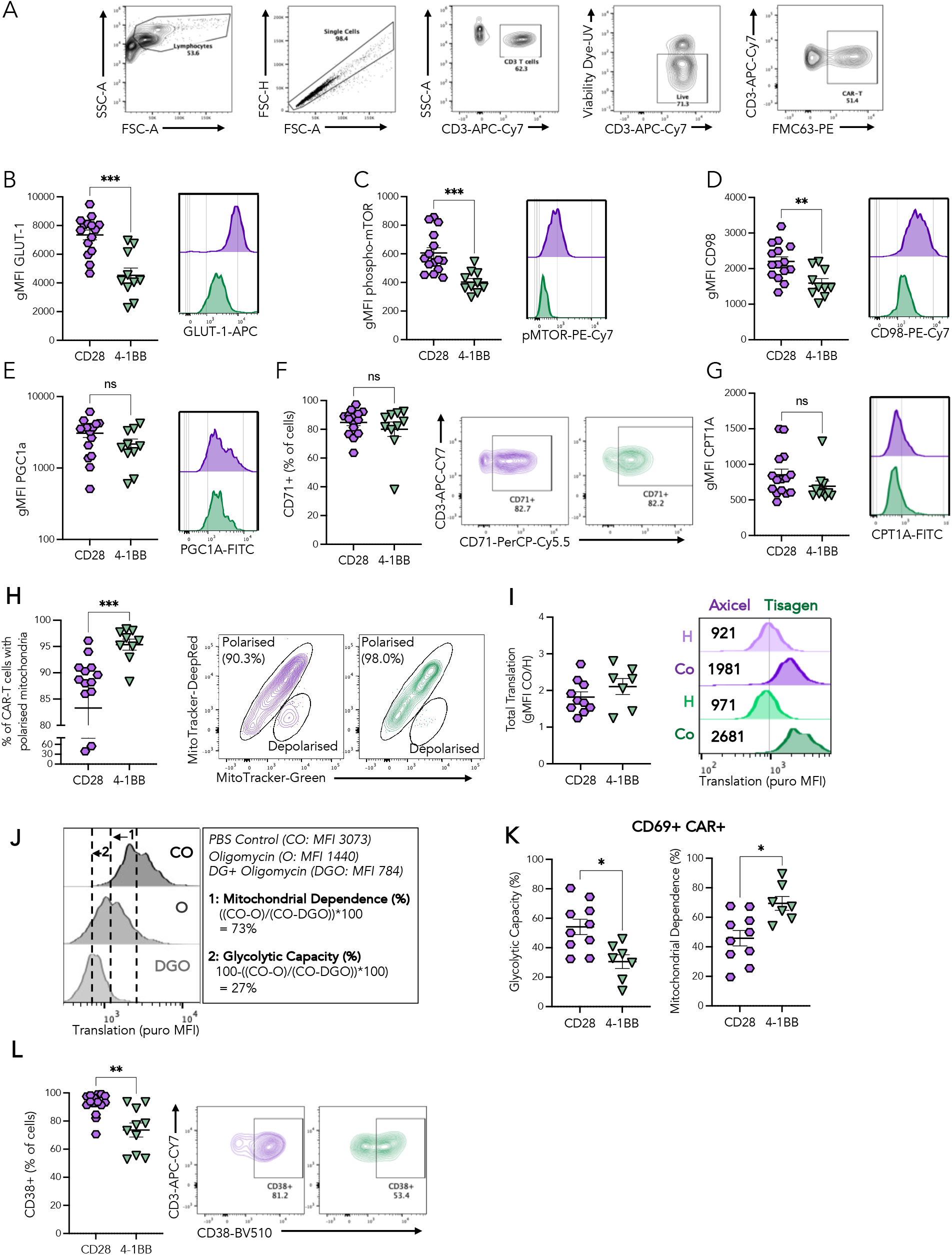
CD28 co-stimulation drives increased expression of glycolysis-associated proteins in patient CAR-T cells. The expression of nutrient transporters and metabolism-associated proteins were assessed by flow cytometry in patient CAR-T cells at day 7 post infusion, comparing patients receiving CAR products with either CD28 or 4-1BB co-stimulation. **A)** Gating strategy for assessing CAR-T cells. **B)** The geometric mean fluorescent intensity (gMFI) of glucose transporter 1 (GLUT-1) **C)** phosphorylated mTOR (p-mTOR) **D)** CD98 and **E)** Peroxisome proliferator-activated receptor gamma coactivator 1-alpha (PGC1a) with representative histograms of fluorescent intensity shown. **F)** The percent of CAR-T cells expressing CD71, with representative gating shown. **G)** The gMFI of Carnitine palmitoyl transferase I alpha (CPT1a). **H)** PBMCs were stained with mitochondrial dyes; MitoTracker DeepRed (polarisation dependent) and Mitotracker Green (polarisation independent). Data shown is the percent of CAR-T cells harbouring polarised mitochondria, which have an equal ratio of DeepRed to Green dyes as shown in the adjacent representative example. **I)** PBMCs were incubated with either PBS (Control; Co) or Harringtonine (H; an inhibitor of protein synthesis) for 45 minutes with puromycin which is incorporated into newly synthesised proteins. PBMCs were then stained with anti-puromycin to assess the amount of translation that had occurred. Representative example of staining shown in CAR-T cells. **J)** PBMCs were incubated with PBS control, Oligomycin (OXPHOS inhibitor) or a combination of Oligomycin and 2-DG (glycolysis inhibitor) before incubation with puromycin. Calculations for determining mitochondrial dependence and glycolytic capacity are outlined. **K)** Summary data showing percent *of* mitochondrial *energy* dependence and glycolytic capacity in CD69+ CAR*-*T cells. **L)** The percent of CAR-T cells expressing CD38, with representative gating shown. Patients treated with CD28 versus 4-1BB CAR-T were compared by Mann Whitney U test, * p <0.05, **p <0.01, *** p<0.001, n.s. not significant.

### Glycolysis fuels protein translation in CD28-co-stimulated CAR-T while 4-1BB co-stimulated CAR-T rely on OXPHOS

Since increased mitochondrial fitness has been linked to 4-1BB signalling^7^, we next investigated mitochondrial membrane polarization in *ex vivo* CAR-T cells. Membrane depolarization is an indicator of electron transport chain dysfunction^20,21^, we find 4-1BB-CAR-T have an increased frequency of T cells harboring functional mitochondria compared to CD28-CAR-T (fig. 1h). To understand if these observations impacted CAR-T cell metabolic pathway usage *ex vivo*, we utilised ‘Single Cell ENergetic metabolism by profiling Translation inHibition’ (SCENIT**H)**. SCENITH relies on protein synthesis being an energetically costly process, such that cellular ATP production correlates with cellular translation levels^11,22^. Translation levels can be measured via incorporation of puromycin into newly synthesis peptides. We found total translation levels to be similar in CD28 and 4-1BB-CAR-T, as detected by puromycin incorporation (fig. 1i). However, the metabolic pathways that fuel translation in the two CAR-T products are distinct. By using the specific inhibitor Oligomycin-A to block ATP synthase in the electron transport chain and 2-Deoxy-D-Glucose to inhibit glycolysis we could delineate metabolic pathway usage (fig. 1j). Assessing the *ex vivo* metabolically active subset of CD69+ CAR-T cells, we find that 4-1BB signalling leads to increased OXPHOS dependence whereas CD28 signalling drives increased dependence on glycolysis (fig. 1k). The NADase ecto-enzyme CD38, which is upregulated on activated T cells, has been shown to impact on mitochondrial function by regulating mitophagy^23–25^. The lower mitochondrial fitness in CD28-CAR-T was accompanied by a significantly increased frequency of CD38+ CAR-T cells as compared to 4-1BB CAR-T (fig1. l). This might cause reduced mitochondrial fitness as a result of impaired clearing of dysfunctional mitochondria due to reduced mitophagy^26^ in CD28-CAR-T. Overall, we find that in patients, CD28 and 4-1BB signalling induce distinct CAR-T metabolic profiles.

### The *ex vivo* CAR-T metabolic profile is associated with cellular differentiation

T cell metabolic pathway usage is associated with T cell functional differentiation status, with glycolysis supporting a more effector-like phenotype and OXPHOS preferentially used by memory T cells^27,28^. We therefore assessed the activation phenotype of patient CAR-T cells. CD69 is an early T cell activation marker and plays an important role in tissue-residence in humans^29^. We find that an increased frequency of CD28-CAR-T express CD69 compared to 4-1BB-CAR-T (fig. 2a). However, CD103, which is co-expressed with CD69 on tissue-resident T cells, was not significantly increased in CD28 versus 4-1BB CAR-T (fig. 2b,c). CD28 CAR-T expressed greater levels of the activation marker and co-inhibitory molecule PD1 compared to 4-1BB CAR-T (fig. 2d). This could suggest greater CAR-T cell activation, as opposed to dysfunction, as the frequency of CAR-T cells expressing CD39, another marker often linked to exhaustion, was not significantly altered (fig. 2e).

**Figure 2.**
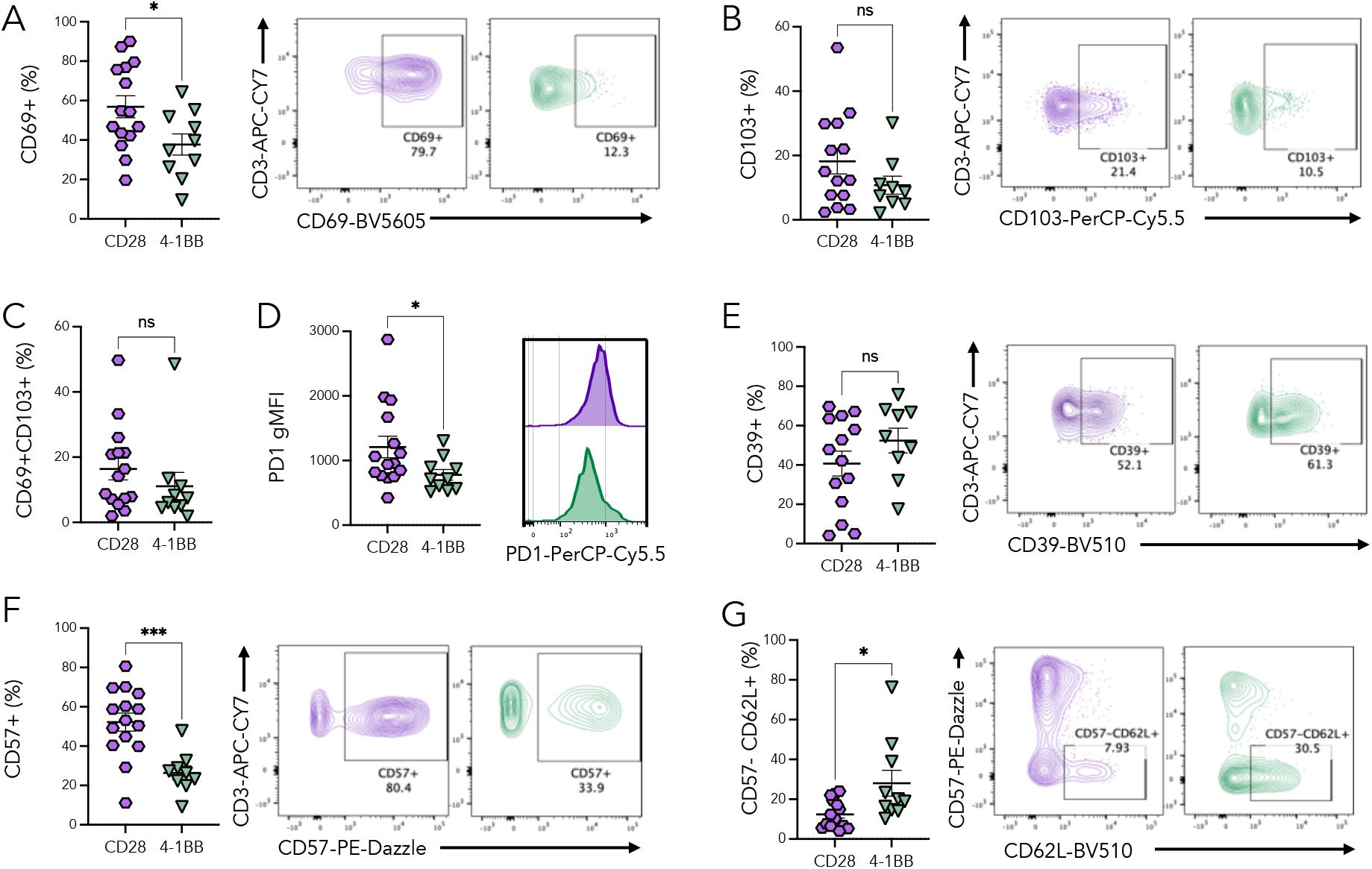
CD28 and 4-1BB co-stimulation drive divergent CAR-T phenotypes. The expression of phenotypic markers was assessed by flow cytometry in patient CAR-T cells at day 7 post infusion, comparing patients receiving CAR products with either CD28 or 4-1BB co-stimulation. **A)** The percent of CAR-T cells expressing CD69 and **B)** CD103 with representative gating shown. **C)** The percent of CAR-T cells co-expressing CD69 and CD103. **D)** The geometric mean fluorescent intensity (gMFI) of programmed-cell-death-1 (PD1) with representative histograms of fluorescent intensity shown. **E)** The percent of CAR-T cells expressing CD39 and **F)** CD57 with representative gating shown. **G)** The percent of patient CAR-T cells that are negative for CD57 expression and positive for CD62L expression, with representative gating shown. Patients treated with CD28 versus 4-1BB CAR-T were compared by Mann Whitney U test, * p <0.05, *** p<0.001, n.s. not significant.

We were interested to assess the expression of CD57, a marker for effector T cell differentiation^30^, which has been previously correlated with successful CAR-T therapeutic outcome^31^. We find a striking increase in the frequency of CD57+ CAR-T cells with CD28 signalling compared to 4-1BB (fig. 2f). Whereas the frequency of CAR-T cells expressing the memory marker L-selectin (CD62L)^32^ was increased by 4-1BB signalling (fig. 2g). Taken together, our *ex vivo* findings confirm the roles of CD28 and 4-1BB signalling in supporting effector and memory CAR-T cell phenotypes respectively in DLBCL patients.

### Increased frequency of circulating CD28 co-stimulated CAR-T likely links to better therapeutic outcome

Having seen clear differences in the metabolism and phenotype of CAR-T cells induced by CD28 and 4-1BB signalling, we next investigated whether this would translate into differential therapeutic response. We first assessed the frequency of CAR+ cells at day 7 post-infusion as an indicator of CAR-T expansion capacity. The frequency of CAR+ T cells has been previously associated with positive outcome in patients^33^. We find a higher frequency of CAR+ T cells in patients who received the CD28-CAR-T compared to those that received the 4-1BB CAR-T (fig. 3a), suggesting increased expansion capacity of CD28-CAR-T. In our limited cohort we see a trend towards increased overall survival (OS) and progression-free survival (PFS) in patients that received CD28-CAR-T (fig. 3b). This is in line with a recent large-cohort study which found significantly improved therapeutic outcomes following CD28 compared to 4-1BB CAR-T therapy^10^. Taken together, our results indicate skewing of CD28-CAR-T towards a glycolytic effector phenotype is accompanied by a higher frequency of circulating CAR-T cells and may link to a better therapeutic outcome in DLBCL.

**Figure 3.**
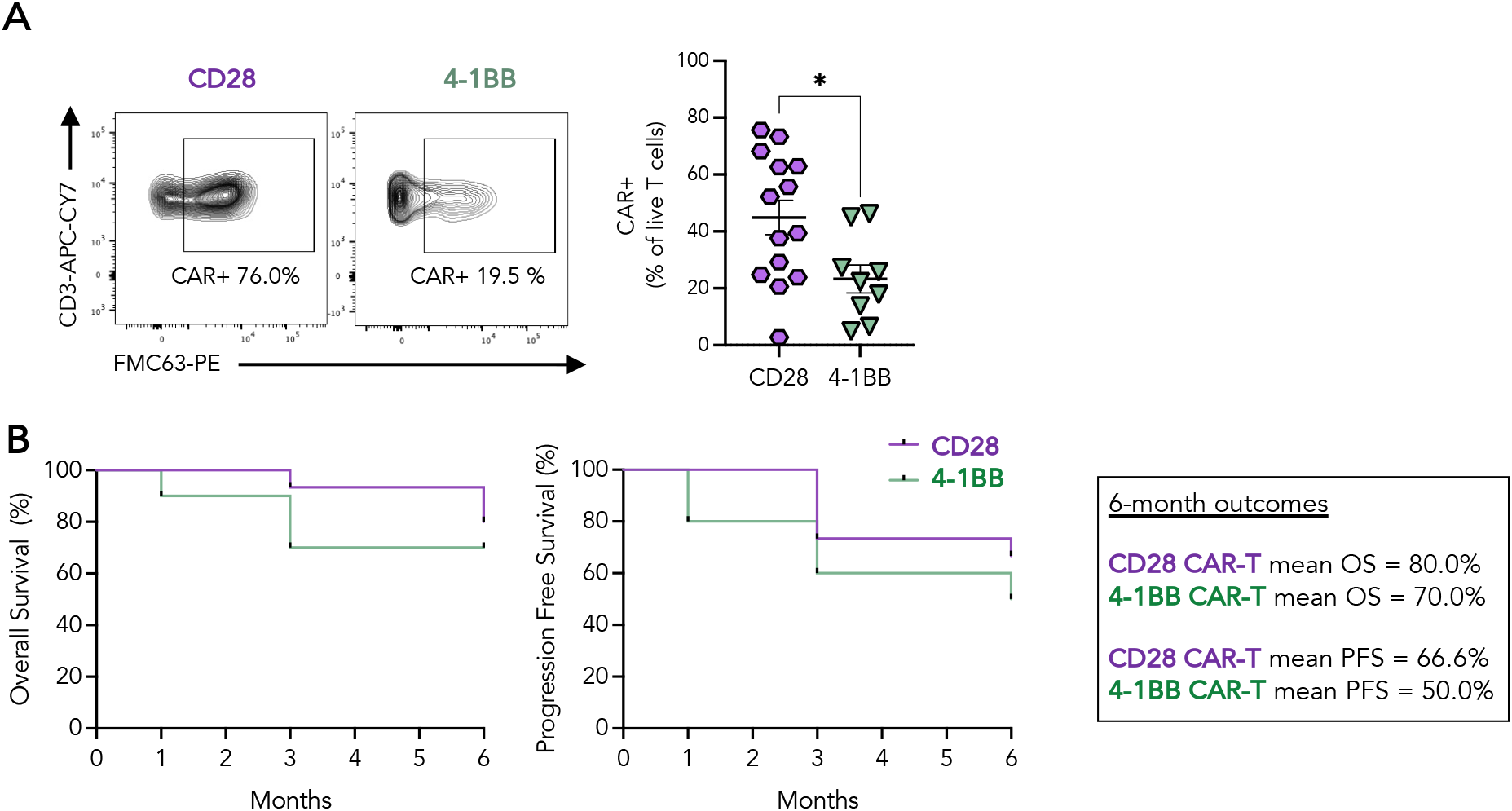
Patient Therapeutic Response to CD28 and 4-1BB CAR-T. **A)** The percent of T cells that are CAR-T, determined by flow cytometry at day 7 post-infusion. **B)** Kaplan-Meier curves of overall survival and progression-free survival in patients treated with CD28 or 4-1BB CAR-T. Patients treated with CD28 versus 4-1BB CAR-T were compared by Mann Whitney U test, * p <0.05.

### A balanced CAR-T cell metabolism correlates with successful clinical response

Since, in our cohort, CD28 co-stimulated CAR-T have a more glycolytic effector-like phenotype, and the CD28 product has been reported to lead to superior clinical efficacy^10^, we next established whether this particular metabolic profile was more prevalent in lymphoma patients who responded to CAR-T therapy. We used PCA analysis to perform unsupervised clustering of patient samples based on all measured CAR-T metabolic and phenotypic parameters. This clustering showed an overall trend for patients to cluster by product received, with CD28-CAR-T (purple) showing separation from 4-1BB (green) (fig 4a). We additionally annotated the data by response at 6 months, with patients being defined as ‘responders’ if they had a complete response or ‘non-responders’ if they had progressive disease, as per the Lugano criteria 2014^34^. PCA analysis revealed greater overlap between CD28 and 4-1BB CAR-T within the responder cohort, whereas within the non-responder cohort these two CAR-T products showed greater separation (fig. 4a). To interrogate these differences further, we stratified our cohort into responders and non-responders and looked at markers that we had found to be different between CD28 and 4-1BB CAR-T. Starting with the metabolic marker, GLUT-1 we found no significant difference in expression within responders, however within non-responders the expression in CD28-CAR-T was significantly higher than in 4-1BB-CAR-T (fig. 4.b). Similarly, the frequency of CAR-T cells with polarised mitochondria was only significantly different within the non-responder cohort (fig. 4c). Equally, we observed a trend towards clearer separation of CD28 and 4-1BB driven phenotype in non-responders versus responders, as demonstrated by CD57 and CD62L expression (fig. 4d,e). Overall, our data suggest that successful CAR-T response requires a more balanced metabolism and phenotype, whilst poor therapeutic outcome is associated with a phenotype that is skewed towards a metabolic extreme.

**Figure 4.**
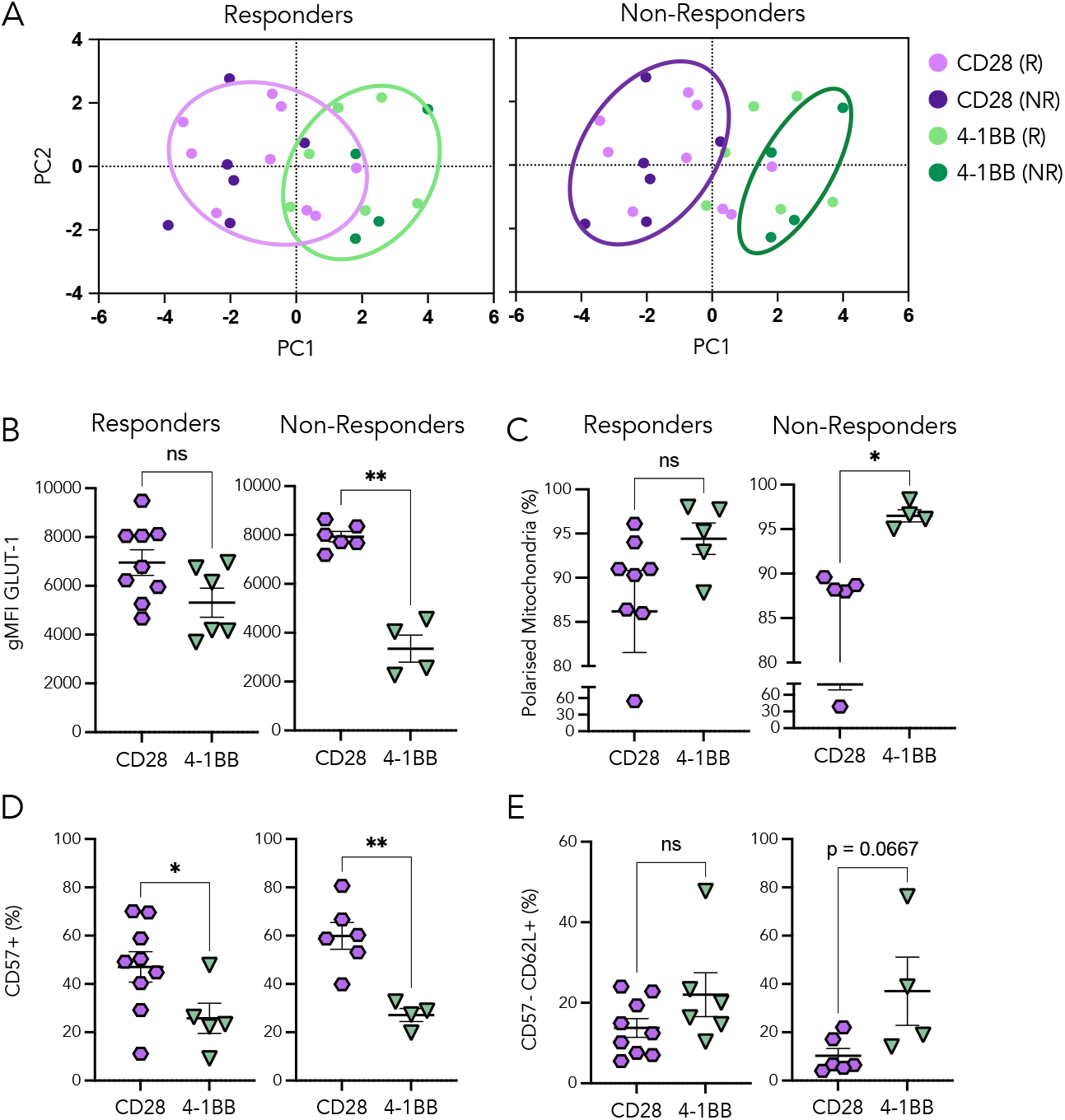
Response to CAR-T therapy is associated with a balanced metabolic profile. **A)** PCA analysis of all assessed metabolic and phenotypic markers in patient CAR-T cells. Patients are categorised by CAR-T co-stimulation, CD28 or 4-1BB, and therapeutic outcome; responders (R) and non-responders (NR) determined at 6 months post-infusion. Plots of the same PCA data is shown side-by-side, with cluster gates applied to responders on the left and non-responders on the right. Metabolic and phenotypic markers were assessed separately in responders and non-responders to CAR-T therapy, comparing CD28 and 4-1BB co-stimulation: **B)** the geometric mean fluorescent intensity (gMFI) of glucose transporter 1 (GLUT-1), **C)** fraction of CAR-T with polarised mitochondria, **D)** fraction of CAR-T expressing CD71, **E)** fraction of CAR-T that are CD57-CD62L+. Groups compared by Mann Whitney U test, * p <0.05, ** p<0.01, n.s. not significant.

## Discussion

We here provide an *ex vivo* study of the metabolic profile of CD19-targeting CAR-T cells derived from patients treated for DLBCL. Although a body of knowledge describing both metabolic and phenotypic features of CD28 and 4-1BB T cell co-stimulation in various *in vitro* and animal models exists, a comparison of these features in CAR-T cells in an *ex vivo* clinical setting remained lacking. In line with previous studies, we found that CD28 co-stimulation led to an effector-like T cell phenotype supported by a glycolytic metabolism, while 4-1BB supported a shift towards increased mitochondrial capacity. Interestingly, molecules phenotypically linked to glycolytic metabolism such as GLUT1 and phosphorylation of mTOR were prominently altered and corresponded well to the functional measure of glycolytic dependence by SCENITH, in contrast classical markers linked to mitochondrial metabolism, such as the transcription factor PGC1a which drives mitochondrial biogenesis^16,18,35^, the iron transporter CD71^19^ and the enzyme CPT1a^36^ were not significantly different between the two different CAR-T cell products. Despite this, when interrogating functional metabolic pathway dependence using SCENITH we found significant reliance of 4-1BB co-stimulated CAR-T cells on mitochondrial OXPHOS, which correlated with a trend towards an increased frequency of CAR-T with polarized mitochondria in the 4-1BB product. Our data thus indicates that in our cohort, GLUT1 is a good predictor of glycolytic dependence, while mitochondrial polarisation could be a predictor for mitochondrial dependence.

Phenotypically the metabolic differences in the two CAR-T products resulted in enhanced effector-like phenotypes in CD28, versus memory-like phenotype in 4-1BB CAR-T. While our study only included a relatively small patient cohort (14 receiving CD28-CAR-T and 9 receiving 4-1BB-CAR-T), patients treated with the CD28 product had a significantly higher frequency of detectable circulating CAR+ T cells 7 days post infusion and we found a non-significant trend for an enhanced clinical response. Although these findings are in line with a large well controlled patient study undertaken by Bachy et al^10^, our clinical findings need to be interpreted with caution, as we were unable to control for patient characteristics such as prior treatment history, ethnicity, sex, age or disease burden. Of note, while similar in all other reported clinical features, patients treated with the 4-1BB-CAR-T product were significantly older than those in the CD28-CAR-T arm and this could potentially have impacted our results (Table 1). The impact of these and further patient characteristics should be taken into account in future studies.

While it is often assumed that a particular metabolic, phenotypic and functional feature will lead to best outcome, our data indicates that a balanced ‘not too much/not too little’ metabolic profile, *combining* metabolic and phenotypic features of effector and memory T cells could be the most successful in driving clinical response in DLBCL. It is likely that a balanced metabolism supports a wider range of effector functions and increased plasticity allows for sufficient persistence, together enabling T cells to mount an effective anti-cancer response. Whether the same is true in other forms of hematological or solid cancers remains to be established. A more comprehensive understanding is needed to allow the production of the most effective CAR-T product for a given disease and patient.

## Materials and Methods

### RESOURCE AVAILABILITY

#### Lead contact

Further information and requests for resources and reagents should be directed to and will be fulfilled by the lead contact, Anna Schurich.

#### Materials availability

This study did not generate new unique reagents.

#### Data and code availability

- This paper does not report original code.
- Data available upon request from the lead contact, Anna Schurich.

### EXPERIMENTAL MODEL AND STUDY PARTICIPANT DETAILS

#### Ethics statement

This study was approved by the King’s College Hospital Local Research Ethics Committee. DLBCL patient samples were obtained from the King’s College London Haematology Biobank which operates under REC approval 23/NE/0160 and HTA License Number 12293. Blood samples from healthy volunteers were obtained under The Guy’s and St Thomas’ License (license number 12121). All subjects gave their informed written consent for inclusion before they participated in the study. The study was conducted in accordance with the Declaration of Helsinki. All storage and record of samples complied with the requirements of the Data Protection Act 1998 and the Human Tissue Act 2004.

#### Patient recruitment and sampling

Patients with DLBCL who received either axi-cel (CD28 CAR-T) or tisa-cel (4-1BB CAR-T) as a treatment at King’s College Hospital were eligible for this study. Patients were recruited between 11^th^ November 2021 and 10^th^ May 2023. Blood samples were taken at 7 days following CAR-T infusion. Samples were processed by the King’s College London Haematology Biobank and isolated PBMCs were stored in liquid nitrogen prior to use. Patient response to treatment was assessed by radiography according to the Lugano criteria (RE**F)**. Peripheral blood was obtained from healthy donors for use in assay optimization and as experiment controls, isolated PBMCs from these samples were stored in liquid nitrogen prior to use.

### METHOD DETAILS

#### CAR-T cell phenotyping

Patient PBMCs were washed with 1xPBS and stained for surface markers with fluorophore-conjugated antibodies in 50 uL 1xPBS for 30 min at 4 °C protected from light. The following antibodies were used for staining: CD3-APC-Cy7, CD8-Alexa Fluor 700, CD69-Brilliant Violet 605, PD-1-PerCP-Cy5.5, CD71-Per-CP-Cy7, CD57-PE-Dazzle CD38-BV510, CD103-PerCP-Cy5.5, CD39-BV510, CD62L-BV510 (all BioLegend), CD98-PE-Vio770 (Miltenyi Biotec). Surface CAR expression was stained for with an FMC63-biotin antibody (CytoArt) followed by a secondary staining with streptavidin-PE (BioLegend) in 1xPBS for 20 min at 4 °C protected from light. Cells were stained concurrently with 1uL/mL blue viability dye (ThermoFisher Scientific). PBMCs were washed with 1x PBS and fixed with fixing reagent (FOXP3 fix/perm kit, ThermoFisher Scientific) for 30 min at room temperature in the dark. PBMCs were washed in permeabilization buffer (FOXP3 fix/perm kit, ThermoFisher Scientific) and stained for intracellular markers using fluorophore-conjugated antibodies in permeabilization buffer for 30 minutes at room temperature in the dark. The following intracellular antibodies were used: Glut1-APC (Abcam), PGC1**α**-FITC (Novus Biologicals) and SER2448-phospho-mTOR-Pe-Cy7 (ThermoFisher Scientific). Samples were acquired on a BD Fortessa. FlowJo v10 software was used for data analysis.

#### Batch Consistency Control

To ensure consistency across batches, we included a healthy-donor control in phenotyping experiments. Prior to experimentation, blood was taken from a single healthy donor, PBMCs were isolated and transduced with CD19-CD28**?** CAR construct. These healthy donor CAR-T were expanded for 10 days in IL-2 and then 40 vials of 2×10^6^ PBMC aliquots were cryopreserved in liquid nitrogen. For every batch of patient cells analysed, a vial of healthy donor control CAR-T cells was analysed in an identical manner. Following acquisition on the flow cytometer, data were normalised to healthy control values using CytoNorm package in R^37^.

#### Mitochondria Polarization Assay

Patient PBMCs were cultured for 16 hours in complete RPMI with 20 IU/mL IL-2. PBMCs were stained with 5nm Mitotracker Deep Red FM and 50nm Mitotracker Green FM (ThermoFisher) in RPMI for 20 minutes at 37°C, 5% CO^2^. PBMCs were washed with ice-cold PBS and stained for surface markers with fluorophore-conjugated antibodies and viability dye in 1x PBS for 30 min at 4°C in the dark prior to flow cytometry analysis.

#### Metabolic flux analysis (SCENITH)

Method adapted from work published by Arguello et al^11^. PBMCs (1 million/mL) were incubated with puromycin (20 μg/mL) and either 1xPBS, oligomycin-A (1μM) or a combination of 2-Deoxy-Glucose (2-D**G)** and Oligomycin-A (1μM) in complete RPMI for 45 min at 37°C, 5% CO^2^. PBMCs were washed in ice-cold PBS and stained for surface markers with fluorophore-conjugated antibodies in 1x PBS for 30 min at 4°C in the dark (table 8.3). PBMCs were stained intracellularly for puromycin with AF488-puromycin antibody (Merck) using the FOXP3 fix/perm kit (ThermoFisher) for 1 hour at room temperature in the dark. Samples were analysed on a BD Fortessa. The following calculations were used to determine mitochondrial dependence and glycolytic capacity as per the published protocol^11^:

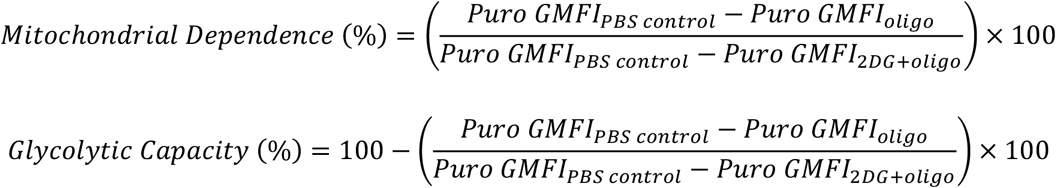

### STATISTICS

Prism 9 (GraphPad) was used for statistical analysis. Patients treated with CD28 versus 4-1BB CAR-T were compared by Mann Whitney U Test. P<0.05 was considered statistically significant.

## Acknowledgements

We sincerely thank all blood donors for supporting our research. We thank the KCH Haematology Biobank for their support. This work was financially supported by: KMRT Studentships to A. Schurich, M. S. Cook; MRC studentship to K. R. Flaherty.

